# Macroscopic cerebral energy efficiency corresponds to neuron reorganization in awake and anesthetized mice

**DOI:** 10.1101/2025.03.08.641356

**Authors:** Da Wang, Hui Li, Yifan Zeng, Jinggui Gao, Mengyang Xu, Binshi Bo, Mengchao Pei, Zhifeng Liang, Ning Zhou, Garth J Thompson

## Abstract

Non-invasive imaging of brain function and energy supply is crucial for diagnosing and treating brain disorders. Conventional imaging struggles to capture altered relationships between energy supply and utilization caused by brain diseases. A novel method, which can be translated to human patients, is to calculate relative power (rPWR) and relative cost (rCST) to assess cerebral energy efficiency. However, whether rPWR/rCST can track individual changes and neural activity remains unproven. Our study compared these non-invasive measures with invasive two-photon microscopy in awake and anesthetized mice. We found that rPWR/rCST distributions were similar between awake mice and humans, but changed in anesthetized mice, indicating a shift in the brain’s economic balance. Furthermore, changes in rPWR/rCST were linked to the reorganization of microscopic neural networks, observed with two-photon microscopy. Our work highlights the potential of rPWR/rCST for medical applications, and that neural network reorganization is linked to the brain’s economic balance.

## Introduction

The brain is one of the most energy-demanding organs, consuming approximately 20% of the body’s total oxygen despite accounting for only 2% of the body’s weight^1^. Extensive research has demonstrated that changes to brain energy consumption are biomarkers for diagnosing neurodegeneration diseases and suggesting treatments^2–4^ and that there is a strong relationship between energy utilization and activity^5–7^.

The development of the brain, both in species evolution and in an individual’s growth, is grounded in the economic balance between minimizing wiring costs and optimizing the patterns of neuronal connectivity ^7–9^. Wiring cost refers to the material and energy expended by the brain to establish and sustain neuronal connections. To reduce these costs, the brain exhibits various topological optimizations, including the utilization of: the shortest paths for neuronal connections^10^; “small-world” networks (i.e., networks with more clusters despite shorter average path length); and other modular structures that facilitate efficient information transmission across different brain regions^11, 12^. Such topological efficiency, which directly impacts brain function, ensures rapid and accurate information transmission, thus enhancing cognitive and processing capabilities^13, 14^. However, changes in physiological states, such as neurodegenerative diseases like Alzheimer’s disrupt the brain’s dynamic network adjustments for cognitive demands, leading to reduced efficiency and functionality due to a shift in the cost-efficiency tradeoff^15^. While the metabolic disruptions in the later stages of such diseases are obvious^16–18^, earlier metabolic changes are subtle, and it is difficult to justify clinical diagnosis at the current low sensitivity levels^18^. Measuring the brain’s energy economics is a promising idea to detect subtle changes early, when intervention may still improve quality of life^19, 20^.

One of the most common methods used to investigate brain energy economics is functional magnetic resonance imaging (fMRI), which measures the blood oxygen level-dependent (BOLD) signal, encompassing various metabolism-related components, including the cerebral metabolic rate of oxygen (CMR_O2_), cerebral blood flow (CBF), and cerebral blood volume (CBV)^21^. Due to its non-invasive nature and high spatiotemporal resolution, fMRI is a potent tool for exploring brain function. Calibrated fMRI methods can provide high-resolution maps of CMR_O2_ from the BOLD signal^22–24^. For example, relaxometry-based calibrated fMRI offers a non-invasive, high-resolution CMR_O2_ mapping technique, utilizing BOLD-sensitive relaxation and CBF data to assess brain activity and metabolism, making it a valuable tool for brain metabolism and activity studies.

Subtle alterations at the microscopic level significantly influence macroscale and whole-brain neurometabolic changes^25^. For example, in experimental models of Alzheimer’s disease, injured neurons in the damaged cortical region can trigger damage to astrocytes^26, 27^, ultimately inducing vascular dysfunction and a decrease in CBF^28^. This, in turn, can significantly impact CMR_O2_^29^. Studying the spatiotemporal correlation between energy metabolism and brain activity is important for connecting macroscopic observations of brain function to the microscopic causes of neurodegenerative disease.

Many studies have directly correlated energy use with brain activity, but neurodegenerative diseases disrupt this relationship^30, 31^. Accurate measurement methods potentially could uncover neurodegenerative disease mechanisms, aiding early diagnosis and treatment. Relative power (rPWR) and relative cost (rCST) have been developed to compare energy supply and demand. rPWR measures relatively how well supply and demand match, thus indexing total neurometabolic activity, whereas rCST measures relatively how much mismatch exists, thus indexing the relative price of neurometabolic activity^32^. rPWR and rCST were derived from both a measurement of the “energy demand”, i.e. neuroglial activity, local functional connectivity density (lFCD) measured by fMRI, and “energy supply,” i.e. a measurement of glucose metabolism, cerebral metabolic rate of glucose (CMR_glc_) measured by ^18^F-fluorodeoxyglucose positron emission tomography (FDG-PET). While this method has proven useful for its generalizability when substituting other measurements of metabolic supply and neuronal activity, and has exhibited sensitivity in detecting alcohol-induced changes in both metabolism and neuronal activity^32^, major questions about this approach remain that are difficult to test in human subjects. First, rPWR and rCST have not yet been proven to track neurometabolic changes between states within individuals. Proving this is critical, as otherwise these measurements may merely be reflecting individual neuroanatomical differences. Second, it is unknown what the changes measured with rPWR and rCST represent in terms of underlying neural activity. These unknowns severely limit the potential of this non-invasive method, as most implantable devices and drugs act on the underlying neural activity or the neurotransmission of individual cells.

To answer these questions, we used a model of awake vs anesthetized mice, combining non-invasive fMRI and invasive two-photon calcium imaging microscopy (2PCI). No previous study has used rPWR and rCST in an animal model, and two-photon imaging cannot typically be used in human subjects, therefore our work uniquely can address questions about the underlying source of rPWR and rCST measurements. We hypothesized that anesthesia would induce consistent alterations in energy utilization efficiency and neuronal network organization. We innovatively employed a relaxometry-based calibrated fMRI technique to derive the CMR_O2_ index as a measure of energy utilization, since CMR_O2_ closely resembles CMR_glc_ in healthy individuals^33, 34^. With the calibrated fMRI method, we were able to obtain rPWR and rCST solely through fMRI, eliminating the need for FDG-PET-associated radiation exposure. Our work demonstrated that the macroscopic balancing of energy supply and demand is intricately linked to the underlying shifts within local microscopic neural networks.

## Materials and Methods

### MRI experiments

#### Animals

This study uses data from the same experimental animals and imaging session as were used in Xu et al. for calibrated fMRI data^35^, Wang et al. for resting-state fMRI data^36^, and both studies for anatomical brain images.

All animal experiments were approved by the Institutional Animal Care and Use Committee of ShanghaiTech University and the Institute of Neuroscience, Chinese Academy of Sciences, China, in accordance with the governmental regulations of China for the care and use of animals.

The preparation of the animals followed the same protocol as our previous work^35^. Briefly, after head surgery, the mice were secured in a head holder and trained to become accustomed to the MRI imaging noise and environment. One week after collecting awake imaging data (Awake), the mice were anesthetized with dexmedetomidine (Dex) to capture images in the anesthesia state. Data from these two sessions (Awake and Dex) for a total of 14 mice were used. The detailed procedures for animal preparation for MRI imaging are described in detail in this previous work.

#### MRI imaging

MRI imaging methods were detailed in our previous work^35, 36^. To summarize, all animals were scanned on a 9.4T MRI system, using a 4-channel cryogenic mouse head coil for transmit and an 86-mm volume coil for receive (Bruker Biospin). Localization scans were done first to center the mouse brain within the imaging field. Different sequences were then used to acquire images, and the full sequence names and details are provided in Table 1. Parameters imaged included the brain structure (T2 Turbo RARE), maps for the T_2_ and T_2_* magnetic relaxation constants (MSME and MGE, respectively), CBF signals (pCASL), and fMRI BOLD signals (EPI) from the mouse brain.

**Table 1:** MRI sequences and their parameters. RARE = rapid acquisition with relaxation enhancement; MGE = multiple gradient echo; MSME = multi slices multi echo; pCASL = pseudocontinuous arterial spin labeling

| Sequence | Echo time<br>(TE) | Repetition<br>time (TR) | Field of view<br>(FOV) | Image size | Number<br>of slices | Slice<br>thickness | Other non-default<br>parameters |
| --- | --- | --- | --- | --- | --- | --- | --- |
| T2 Turbo RARE | 334ms | 3300ms | 16×16mm<sup>2</sup> | 256×256 | 20 | 0.5mm | Averages: 2 |
| MGE | 2.15-17.23ms | 500ms | 22×16mm<sup>2</sup> | 110×80 | 20 | 0.5mm | Echoes: 8, averages: 4 |
| MSME | 10-100ms | 3000ms | 22×16mm<sup>2</sup> | 110×80 | 20 | 0.5mm | Echoes: 10, average: 1 |
| pCASL | 22ms | 4000ms | 71×71mm<sup>2</sup> | 15.975×15.975 | 20 | 0.5mm | Label duration: 3000ms<br>Post labeling delay: 600ms<br>Pairs of label/control: 80<br>Inversion efficiency: 0.59 |
| EPI | 15ms | 1000ms | 15.975×12.15mm<sup>2</sup> | 71×54 | 20 | 0.5mm | Repetitions: 600 |

### fMRI processing method

fMRI processing methods were detailed in our previous work ^35, 36^. In summary, the fMRI BOLD data underwent preprocessing and analysis using custom MATLAB scripts, including motion correction, slice timing adjustment, anatomical image registration to the mouse brain template, spatial smoothing, and filtering of voxel time series.

For noise removal, epochs with low signal-to-noise ratios for brain networks were excluded, and group independent component analysis was used to remove noise components and enhance the quality of the BOLD signal. As the primary somatosensory cortex of the forelimb (S1FL) exhibits a strong correlation between the left and right hemispheres, lacking correlation in S1FL is an indicator of disrupted brain networks in general, potentially signifying an underlying physiological issue. Thus, data with Pearson’s correlation coefficient (PCC) less than 0.3 between the bilateral S1FL were excluded as several previous studies have done ^37–39^ To reduce noise and improve the quality of the BOLD signal, group independent component analysis (ICA) was conducted using GIFT toolbox (http://mialab.mrn.org/software/gift/) as several previous studies have done ^36, 40^. The signal from ventricles and white matter, components 1, 2, 4, and 8 were removed (Supplemental Fig. 5).

### Local FCD estimation

Functional connectivity density mapping (FCD) mapping is a voxel-wise graph theoretical approach that aims to identify hubs within the brain’s networks. FCD involves counting the number of local or global functional connections for each voxel in the brain, utilizing Pearson’s correlation coefficient as a measure of connectivity^41^. Following Shokri-Kojori et al., here local FCD (lFCD) was used to calculate the number of voxels linked with nearby voxels. In accordance with previous work in mice^36^, the threshold for Pearson’s correlation coefficient (PCC) was set to *r* ≥ 0.25, and the nearby cluster size was set at a radius of 2 mm^42^.

### Removal of signals related to temporal signal to noise variation

As per the methods of Shokri-Kojori et al., after smoothing and removing the signal from ventricles and white matter as described above, the voxel-wise temporal signal to noise (tSNR) of fMRI data were calculated using the mean to standard deviation ratio of the raw fMRI time series^32^. Here, rPWR and rCST were calculated by FCD and CMR_O2_, to improve the quality of rPWR and rCST, we calculated the correlation coefficient between tSNR and lFCD, CMR_O2_, rPWR, and rCST. The brain regions that had a significant correlation with rPWR and rCST as shown in Supplemental Tables 3 and 4. These regions were removed for further analysis and are shown in black on the brain maps in Figure 1.

**Fig. 1.**
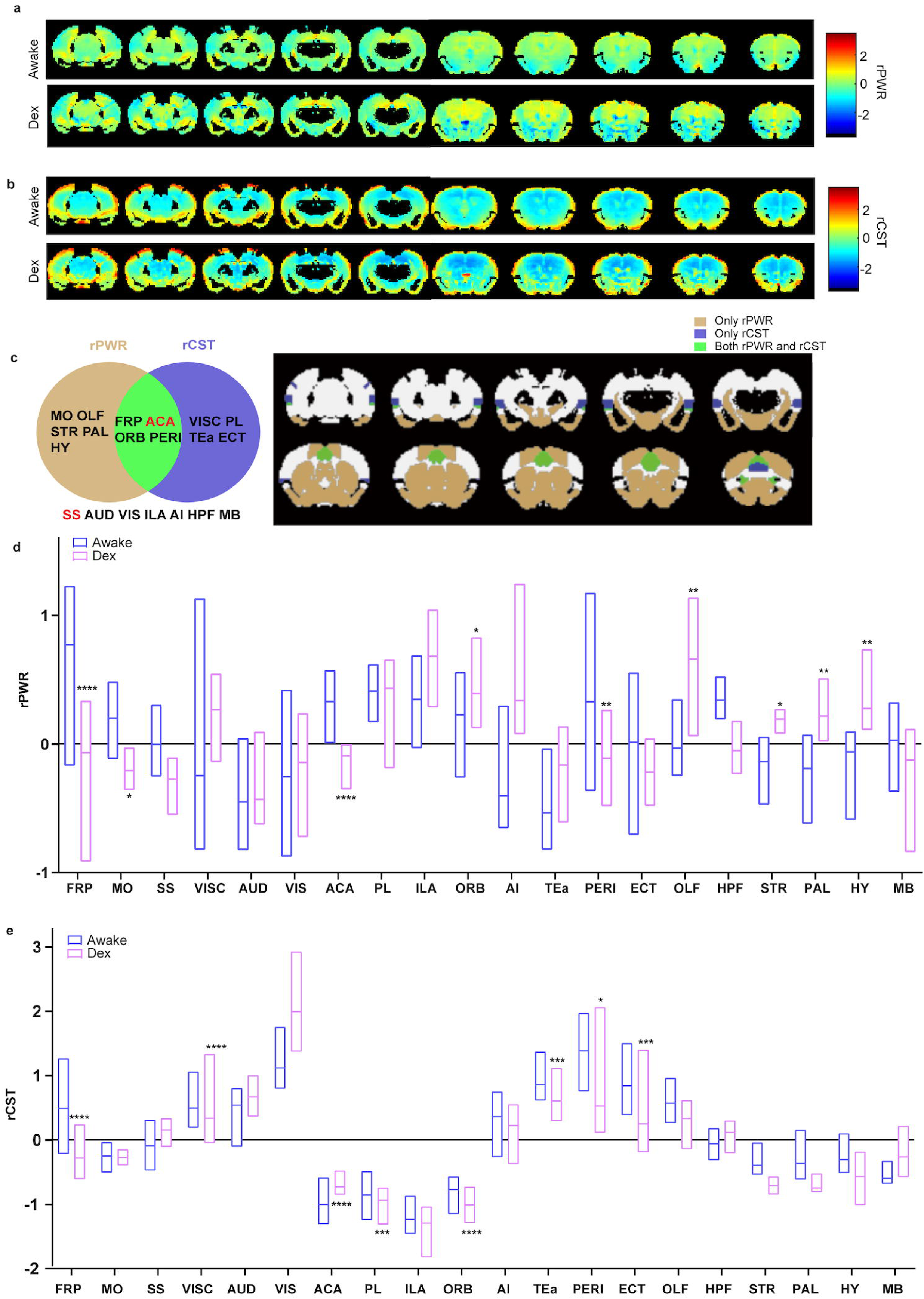
rPWR and rCST under Awake and Dex conditions. **a**, **b** Average rPWR (a) and rCST (b) across all mice in awake and Dex groups in 10 coronal slices, from caudal to rostral (n = 14 mice). Brain regions where CMRO2 or lFCD significantly correlated with tSNR (see Materials and methods) were removed and are depicted as black. Each scatter plot displays data from one mouse. **c** Summary of rPWR and rCST statistical results. The brown and blue circles indicate brain regions that exhibit statistically significant differences between Awake and Dex in rPWR and rCST, respectively. The intersection of these two circles indicates areas that exhibit statistically significant differences in both rPWR and rCST, and regions below the circles exhibited no statistically significant differences in either rPWR or rCST. **d, e** Comparison of rPWR (d) and rCST (e) averaged for 20 brain regions. Data shows the median maximum and minimum values (paired-t test, detailed p value and t value information is shown in Supplemental Table 2. * indicates p < 0.05, ** indicates p < 0.01, *** indicates p < 0.001,**** indicates p < 0.0001). FRP = frontal pole; MO = somatomotor areas; SS = somatosensory areas; VISC = visceral areas; AUD = auditory areas; VIS = visual areas; ACA = anterior cingulate area; PL = prelimbic area; ILA = infralimbic area; ORB = orbital area; AI = agranular insular area; TEa = temporal association areas; PERI = perirhinal area; ECT = ectorhinal area; OLF = olfactory area; HPF = hippocampus formation; STR = striatum; PAL = pallidum; HY = hypothalamus; MB = midbrain.

### Calibrated fMRI

The relaxometry-based calibrated fMRI method was used to obtain relative CMR_O2_, shown in equation (E1). This method was used as it does not require a gas challenge ^35^. The value of *A* was determined by referencing the awake state CMR_O2_ reported previously of 116 μmol/100g/min^43^. Using the average of R_2_’ and CBF measurements from awake mice and equation (E1), we calculated *A* = 69.3257. Note that this *A* value was only used as an estimate to allow per-subject calculation of rCMR_O2_, we do not claim that values produced using this *A* value are quantitative CMR_O2_. In accordance with previous work in mice, we used *ɑ* and *ß* of 0.2 and 0.8, respectively (The *ß* used was the average of the *ß* brain map)^35^.

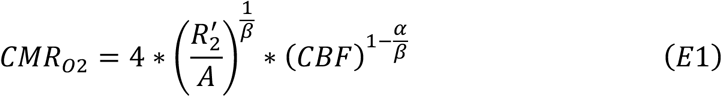

While rPWR and rCST were estimated following Shokri-Kojori et al. (https://github.com/eshoko/COMET)^32^, we used CMR_O2_ instead of CMR_glc_. This replacement was possible because Shokri-Kojori obtained similar rPWR and rCST results with different metabolic supply measurements (CBF) and since the oxygen-glucose index is similar across the brain of healthy individuals whether oxygen is measured with PET or with Calibrated fMRI^44, 45^. As described above, the CMR_O2_ of certain brain regions correlated with temporal signal to noise and were removed.

### Two-photon calcium imaging experiments

#### Animals

Two-photon imaging was conducted only after results were known for rPWR and rCST. We selected one brain region where rPWR and rCST significantly changed between Awake and Dex groups and one region where changes were not significant. In choosing these regions, we also considered the degree of change, looking for areas with large changes (i.e., significant) and areas with small changes (i.e., non-significant) as well as the feasibility of imaging these regions with two-photon imaging.

Four male C57BL/6J mice (2 mice per each chosen brain region) were purchased from the Animal Core Facility of the corresponding authors’ institution. All animal experiments were performed in accordance with procedures approved by the Institutional Animal Care and Use Committee of the corresponding authors’ institution and were consistent with local governmental regulations for the care and use of animals. All mice were housed in strictly controlled environments, maintaining a humidity level of 50 ± 10% and a temperature range of 20 to 26 °C. Mice were exposed to a regular 12-hour light/12-hour dark cycle and had unrestricted access to rodent chow and water. Prior to commencing the experiments, all mice were quarantined for a week to ensure they were free of specific pathogens.

#### Two-photon calcium imaging

For the two-photon calcium imaging (2PCI), a precision micro handheld skull drill was employed to fashion a 4 mm-diameter cranial window precisely positioned above the virus injection site or its symmetrical counterpart relative to the sagittal suture. Following this, a glass slide measuring 4 mm in diameter and 100 μm in thickness, along with a drug administration cannula, were gently inserted into the cranial window and securely adhered using biological tissue glue. After a recovery period of at least one week, an imaging baseplate was meticulously placed on the cranial window and firmly adhered with dental acrylic resin. The miniature two-photon microscope (FHIRM-TPM V2.0, Transcend Vivoscope Biotech), although detachable, features a bracket that is permanently affixed to the baseplate situated above the cranial window. Prior to imaging, both the bracket cover and the protective gel covering the cranial window were carefully removed. Subsequently, the headset was seamlessly mounted onto the bracket and securely locked in place using an M2 screw, ensuring stability and precision during the imaging process.

Mice were injected with AAV2/9-hsyn-GCaMP6f-WPRE-pA. After recovery from virus injection, mice were trained for awake 2PCI experiments (Supplemental Fig. 6a). The emission filter of GCaMP6f has a wavelength range of 500 to 550 nm and an excitation wavelength of 920 nm. Image data were acquired using Imaging software (GINKGO-MTPM, Transcend Vivoscope Biotech) at a frame rate of 9.76Hz, total imaging time was 10 minutes. Two full sessions were recorded for each mouse and each condition. The Field of view (FOV) size was 420 × 420 μm (512 × 512 pixels). The average power delivered to the brain was less than 110 megawatts. After each recording, the focal plane and imaging position are checked and manually re-aligned with the initial image if necessary. Upon acquiring the calcium fluorescence data, immunofluorescence staining was subsequently performed to confirm the presence of viral injection in ACA (Supplemental Fig. 6b), and in the SS (Supplemental Fig. 6c).

### 2PCI processing

Somatic regions were identified by STNeuroNet (https://github.com/soltanianzadeh/STNeuroNet), and manual calibration in ImageJ (https://imagej.net/software/imagej/).

Calcium imaging data were analyzed with custom-written MATLAB software. The fluorescence signal of each neuron was extracted from identified neurons using the annular ring subtraction (ARS) algorithm^46^, and the fluorescence change relative to the baseline (‘ΔF/F’) was calculated as the calcium transients. The frequency of calcium events was quantified as the number of occurrences per cell per minute. The amplitude of calcium transients was measured by the change in fluorescence (ΔF/F), and the duration was characterized by the full width at half maximum (FWHM) amplitude. All of these values were calculated using custom-written MATLAB scripts.

### Small-worldness calculation

Small-worldness (SW) was characterized by high global cluster coefficient *CC* and short average path length *L* (E2)^47^. Cluster coefficient measured the degree to which nodes in the network tend to cluster together (E3)^47^. For a node *i*, the local clustering coefficient *CC_i_* is the ratio of the number of edges *E_i_* between the node’s neighbors to the maximum possible number of edges between them. *k_i_* is the number of connections between node *i* and other nodes. *CC* is global cluster coefficient, is the average of *CC_i_*. The average path length is the mean of shortest path lengths between all pairs of nodes in the network (E4). *N* is the number of nodes. *N_hj_*(*i*) is the shortest number of paths between node *j* and node *h* through node *i*, *n_hj_* is the shortest number of paths between node *j* and node *h*.

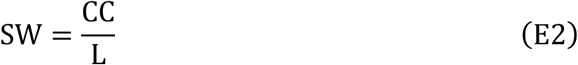

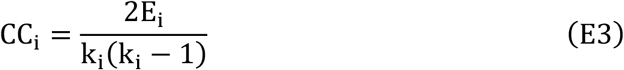

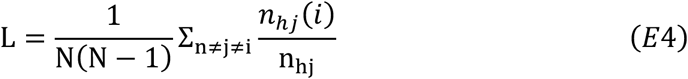

### Statistical analysis

Paired *t* tests were performed, examining the effect of anesthesia on metrics derived from MRI and 2PCI data. The metrics upon which significance testing was conducted were rPWR, rCST, lFCD and CMR_O2_ from MRI data, and frequency, amplitude, duration, and small-worldness from 2PCI data. The *p* values were corrected based on the sequential goodness of fit metatest for multiple comparsions^48^.

## Results

### Relative power (rPWR) and relative cost (rCST) derived from functional connectivity density (lFCD) and cerebral metabolic rate of oxygen (CMR_O2_)

In our study, we adopted an approach that uses CMR_O2_ as a surrogate for the CMR_glc_ that was employed previously^32^. Prior research has indicated that, in healthy subjects, CMR_O2_ and CMR_glc_ exhibit comparable patterns^33, 34^, suggesting that it is reasonable to employ CMR_O2_ as an indicator of energy supply (Supplemental Fig. 1). Also, by using the calibrated fMRI method, we were able to obtain rPWR and rCST using only MRI (Supplemental Fig. 1). Here we build on our previous work with mouse models comparing the awake state with anesthetized conditions, a state change which is highly useful for elucidating the intricate relationship between energy supply and metabolism^35, 36, 49^. Numerous prior studies have demonstrated that anesthesia significantly modulates the tricarboxylic acid (TCA) cycle^50, 51^, subsequently inducing alterations in CMR_O2_ and CBF^52^. Leveraging the mouse as our primary research subject, we were later able to further probe the neuronal-level changes induced by various anesthetic states, utilizing invasive optical imaging technology.

### rPWR and rCST changes between the Awake and Dex groups within the same mice

We recorded fMRI, to measure the energy demand using lFCD, and calibrated fMRI, to measure the energy supply using CMR_O2_, from which we calculated rPWR and rCST in two states, awake mice (Awake) and the same mice anesthetized with dexmedetomidine (Dex). A map of the whole brain and a bar plot of average results in 20 brain regions is shown for rPWR and rCST (Fig. 1a, b). Our results showed that rPWR and rCST had significant, region-specific state changes between the Awake and Dex states within the same mice (Fig. 1c-e). The trends of changes in rPWR and rCST due to anesthesia effects exhibited a distinct pattern. For example, In the anterior cingulate area (ACA) brain region, there was a significant increase in rPWR and a concurrent significant decrease in rCST (*p* = 1.19 × 10^−7^ *T* = 7.22 rPWR, *p* = 8.29 × 10^−4^ *T* = −3.78 rCST). Previous studies have shown that CMR ^35^ and lFCD^36^ were lower in Dex as compared to Awake for almost all brain regions (Supplemental Fig. 2). In contrast, the change we observed in rPWR and rCST was very heterogenous (Fig. 1d, e).

### Comparison of rPWR and rCST between humans and awake mice

While Shokri-Kojori, et al. focused on human populations^32^, our study marks the first application of rPWR and rCST in a mouse model. Healthy human data showed higher rPWR in the default model network (DMN), which indexes high activity and energy demand. This is consistent with the DMN network being the most active and energy demanding network^53–55^. Similar to human data, our awake mouse data showed higher rPWR in the ACA region (Table 2 and Supplemental Fig. 3a), which, in rodents, is believed to have homology with the human DMN^56^. The highest rPWR between the 20 brain regions was the FRP region, which can be explained by a high concentration of proteins related to the tricarboxylic acid cycle in the FRP region^57^. Lower rCST was also found in the ACA for both humans and mice (Table 2 and Supplemental Fig. 3b).

**Table 2:**
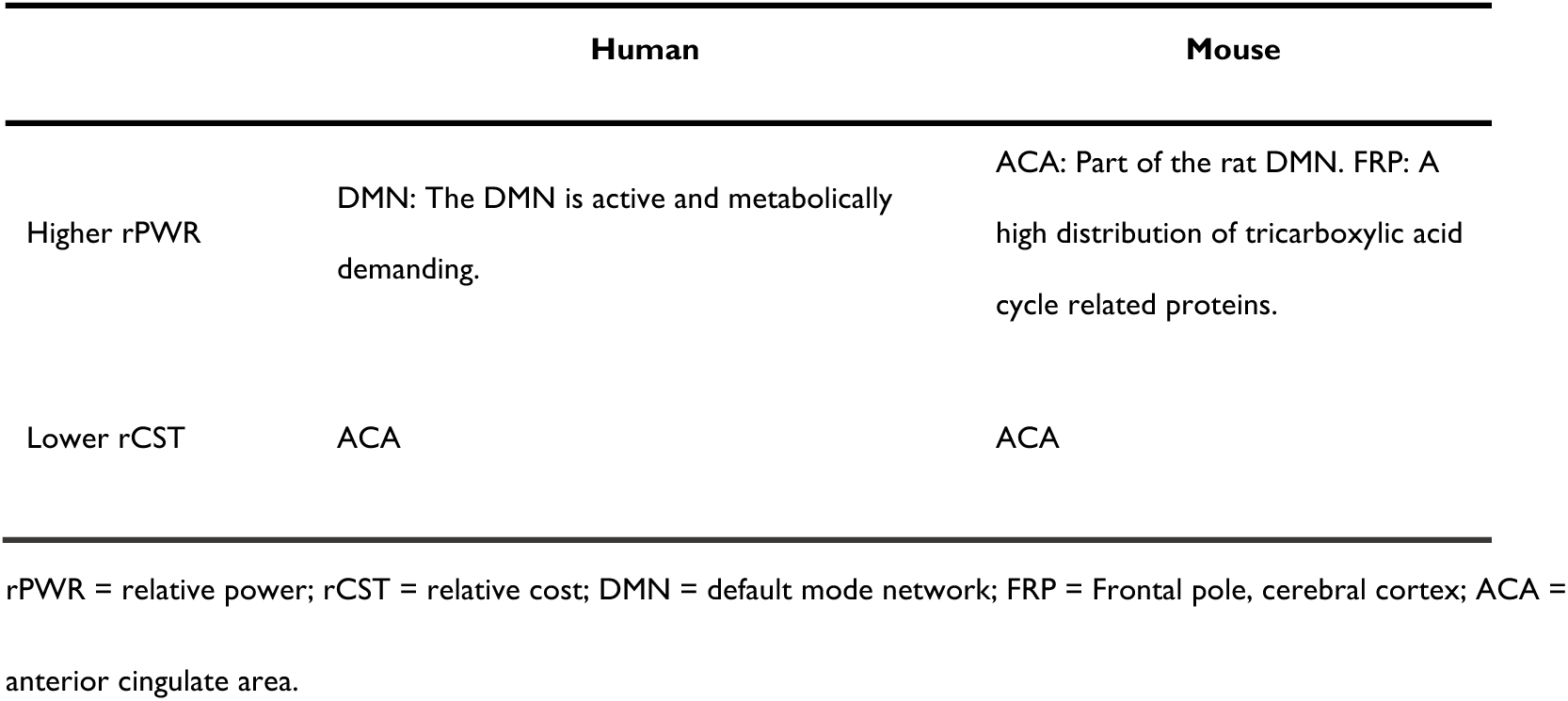
Comparison of rPWR and rCST between awake humans and awake mice.

### Enhanced Differentiation of ACA and SS regions through rPWR and rCST metrics

rPWR and rCST demonstrate a more pronounced divergence in per-brain region changes compared to the lFCD and CMR_O2_. When employing FCD or CMR_O2_, the anterior cingulate area (ACA) and the somatosensory (SS) regions exhibit similar patterns, lacking any significant deviation from the overall dataset (Fig. 2a, b)^35, 36^. However, upon the application of rPWR and rCST, a stark contrast emerges between the ACA and SS regions (Fig. 2c, d). The ACA, in particular, emerges as an outlier with statistically significant differences, highlighting its distinct response to anesthesia. This observation demonstrates that unique insights are provided by rPWR and rCST, metrics that have not been previously utilized to assess state changes in brain activity. Our findings suggest that these relative metrics offer a more nuanced understanding of brain region dynamics, potentially unveiling distinct neurophysiological signatures that were not apparent with the original metrics.

**Fig. 2.**
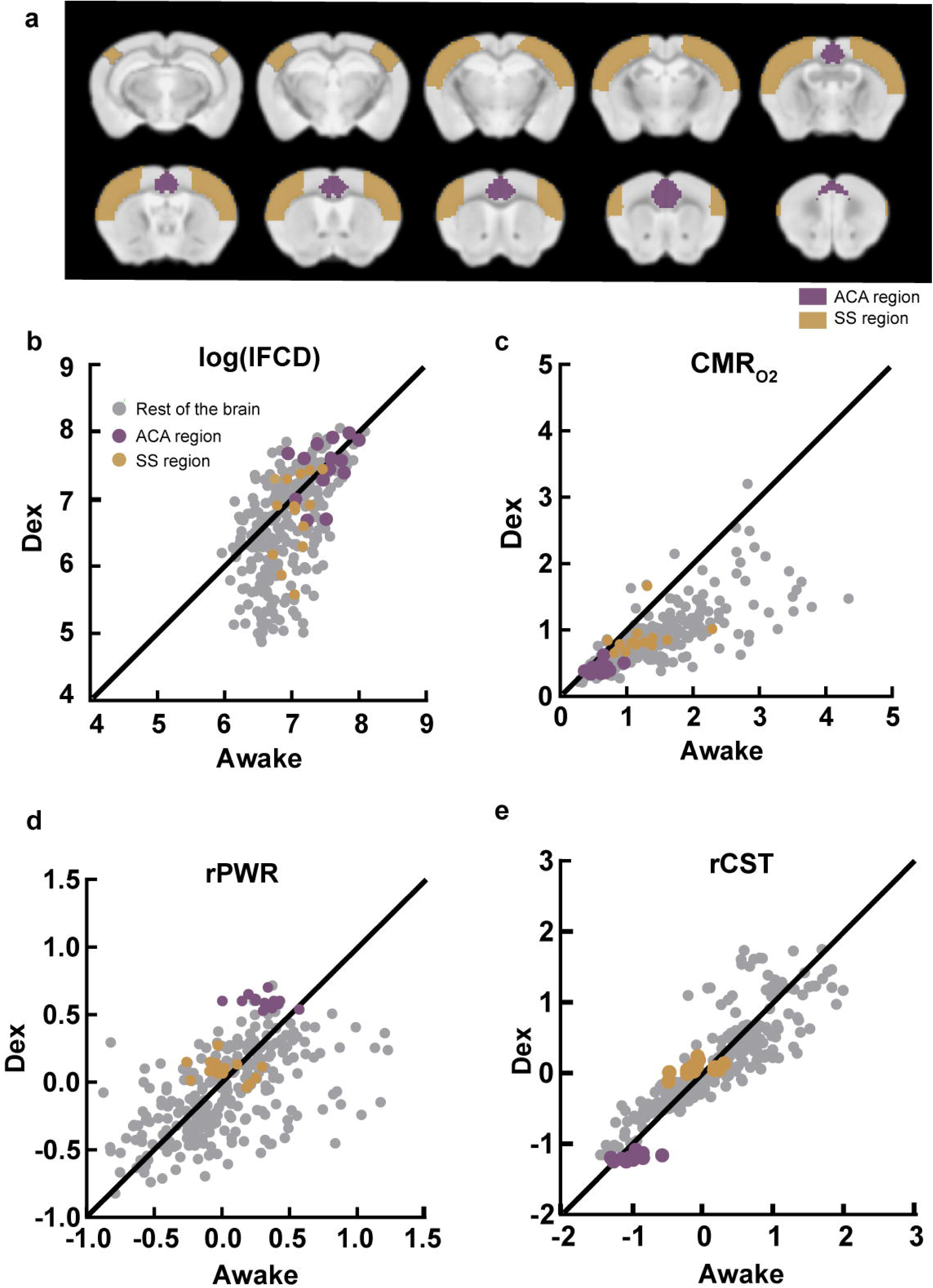
Scatter plots of natural logarithm of lFCD, CMRO2, rPWR and rCST. **a** Brain map (as per Fig. 1a, b, c) showing the locations of ACA region and SS regions. **d-e** Each dot is one mouse’s value in one brain region (n = 14 mice). Purple, yellow and gray circles are values from the ACA, SS, and remaining brain regions. Above the diagonal indicates a higher value under anesthesia, and below indicates a higher value when awake. The values on the diagonal are where awake and anesthesia states are approximately equal. lFCD = local functional connectivity density; CMR_O2_ = cerebral metabolic rate of oxygen; rPWR = relative power; rCST = relative cost; ACA = anterior cingulate area; SS = somatosensory areas.

### Anesthesia induced a decrease of neuronal activity within the SS and ACA regions

Based on our rPWR and rCST results, we selected the SS region as a region which showed no or minimal response to anesthesia, and the ACA as a region which showed a strong and statistically significant response to anesthesia. We performed 2PCI experiments under Awake and Dex conditions in mice with the imaging field of view located in the SS region and, in a separate cohort of mice, in the ACA region.

Data from 2PCI experiments was analyzed to examine changes in the neuronal activity caused by anesthesia (Fig. 3a, b). We measured the frequency, amplitude and duration of calcium transients (Fig. 3c-h). We found that within the SS brain region, the frequency and amplitude of calcium transients was significantly decreased under Dex (*p* = 0.011, *T* = −5.61 from frequency, *p* = 0.024, *T* = −4.23 from amplitude). The duration of calcium transients did non-significantly change under Dex. Within the ACA brain region, there was no significant difference for either amplitude or duration between Awake and Dex, and the frequency was significantly decreased under Dex (*p* = 0.016, *T* = −4.94).

**Fig. 3:**
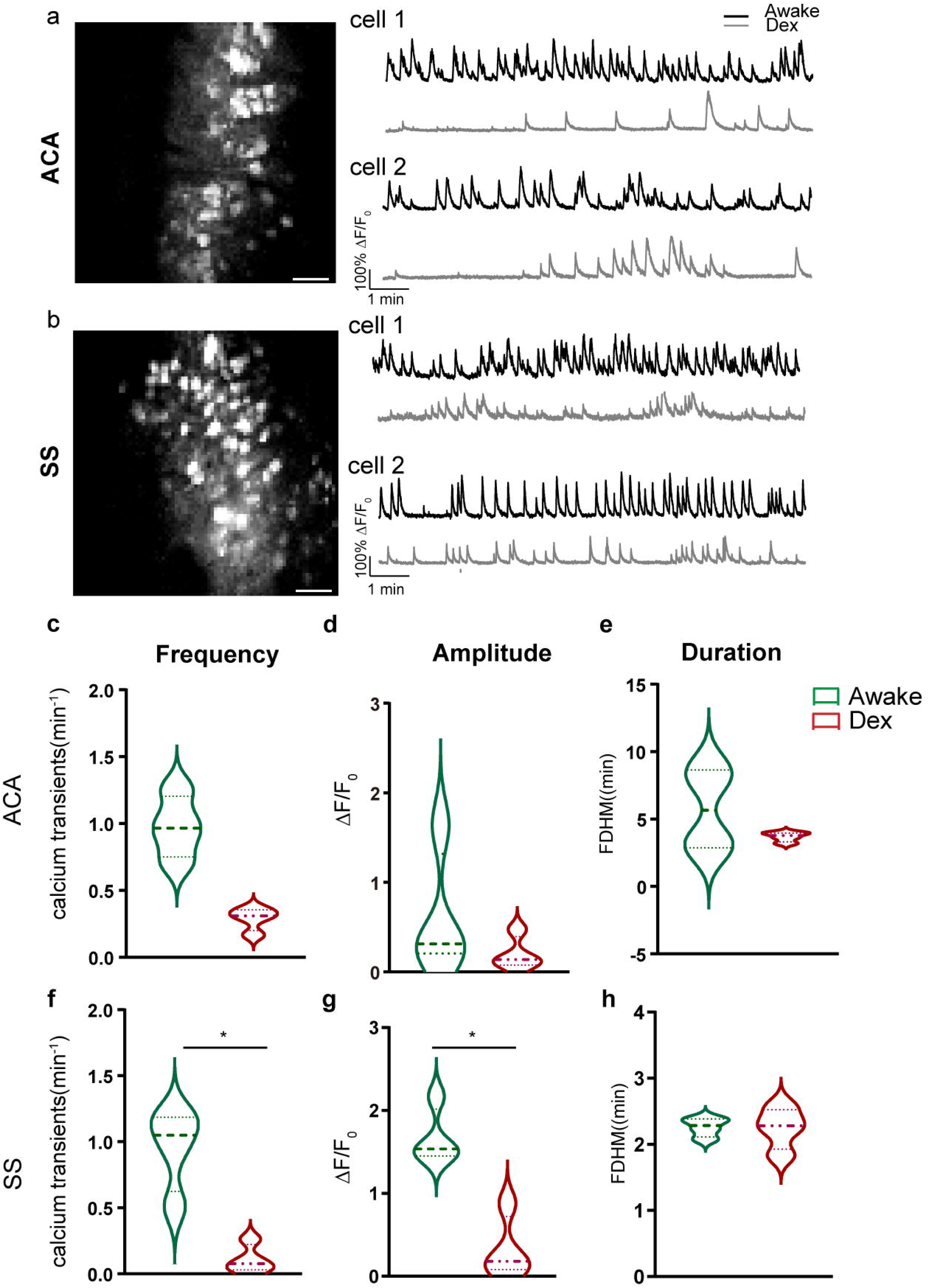
Calcium transients within ACA and SS regions under Awake and Dex conditions. **a**, **b** Single cell calcium intensity within the ACA (a) and SS (b) brain regions under different anesthesia states. **c-h** Frequency, amplitude, and duration of calcium events within the ACA and SS brain regions under the different states (n = 4 mice, 2 per region, each mouse collected twice). The upper and lower dotted lines represent the third and first quartiles, respectively, and the thick dotted line is the median. Asterisks (*) indicate statistically significant differences (p < 0.05) between the awake and anesthetized states in the corresponding brain regions for the corresponding metrics. ACA = anterior cingulate area; SS = somatosensory areas; FDHM = full duration at half maximum amplitude.

### Small-worldness increased under Dex within the ACA region only

Graph theoretical methods are highly effective in analyzing neuronal networks because each neuron can be treated as a node, with the communication between them measured to assign edges based on their degree of connectivity. In particular, small-worldness measurements are widely used to assess the structure and function of networks in biological studies^58, 59^.

A heatmap depicting the Pearson’s correlation coefficient (PCC) between calcium fluorescence signal time series from 10 representative neurons in a single mouse is shown (Fig. 4a, d). Examining PCC across all mice and all neurons, we found no significant change of PCC between Awake and Dex groups in either brain region (Fig. 4b, e). Subsequently, we calculated the small-worldness based on these PCC values. Small-worldness was calculated by cluster coefficient and average path length, however, neither of these individual measurements displayed a significant change between Awake and Dex groups in neither the ACA nor SS regions (Supplemental Fig. 4). Our findings revealed a significant decrease of small-worldness within the ACA region under Dex (*p* = 0.0399, *T* = 3.485) (Fig. 4c). In contrast, no significant change of small-worldness was observed within SS region (Fig. 4f). However, despite this difference in network structure, significant changes in calcium transients were similar for these two brain regions (Fig. 3c-h). Our results show that, compared to the SS brain region, the neural network in the ACA brain region was altered, likely to maintain neuronal activity under Dex. This enhancement of the small-worldness in the ACA brain region under anesthesia indicates a reconstruction of the neural network, making the neural network more economically efficient^60^.

**Fig. 4:**
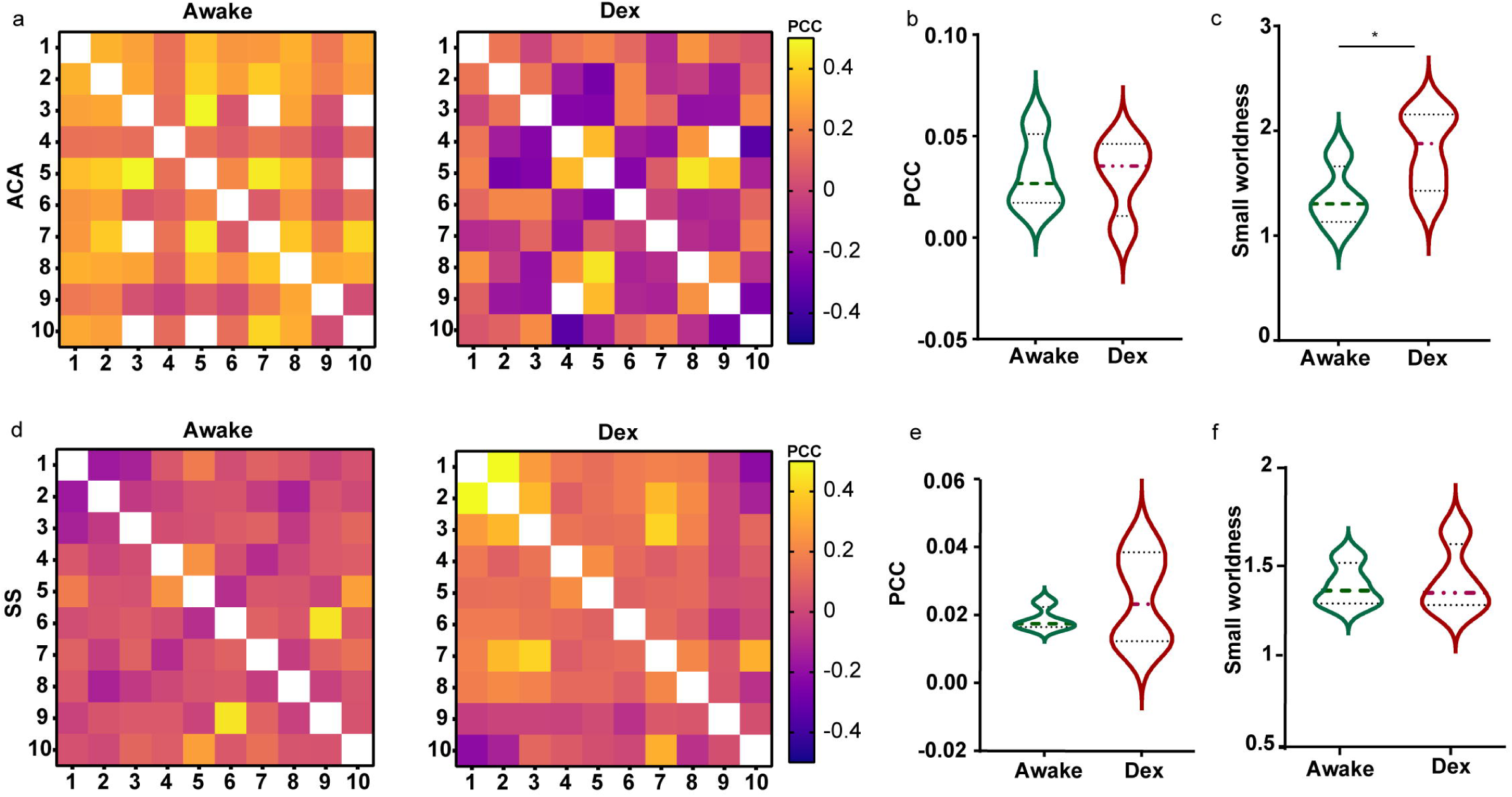
Pearson correlation coefficients of calcium signals in time series of 10 neurons and changes of small-world networks between awake and anesthesia (Dex) states. **a**, **d** Heat map of PCC value of the calcium signal from the ACA and SS regions, with the white squares indicating a 1.0. Each row and its respective column are one representative neuron. All neurons for a given region are from the same mouse. **b, e** PCC under Awake and Dex conditions from ACA and SS brain regions (n = 4 mice, 2 per region, each mouse collected twice). The upper and lower dotted lines represent the third and first quartiles, respectively, and the thick dotted line in the middle is the median. There was no significant difference in PCC under Awake and Dex conditions in either brain region. **c, f** Small-worldness under Awake and Dex conditions from ACA and SS brain regions, marked as in b, e (n = 4 mice, 2 per region, each mouse collected twice). Small-worldness was significantly higher in Dex group compared with the Awake group in the ACA brain region only (paired-*t* test, *p* = 0.0399, *t* = 3.485). In the SS brain region, there was no significant difference between Awake and Dex groups, or even a clear trend. PCC = pearson’s correlation coefficient; ACA = anterior cingulate area; SS = somatosensory areas; Dex = anesthetized by dexmedetomidine;

### Correspondence between rPWR/rCST and the underlying neural networks

Fig. 3 demonstrates that within the ACA, where rPWR and rCST significantly changed, calcium transients were similar to the SS, a region where rPWR and rCST did not significantly change. This suggests that despite reduced neuronal activity, the ACA network adapts to maintain essential functions. We observed that lower rCST levels correlated with neural network efficiency (Fig. 4), allowing functional integrity under energy constraints. Correspondingly, the ACA brain region is closely related to a series of critical high-energy functions, including but not limited to reward processing, decision-making, and emotional responses^61^.

Our findings establish a correspondence between rPWR/rCST, which characterize the match/mismatch between energy supply and neuronal activity, and properties of the underlying neural networks. The increase in macro-level energy utilization efficiency reflected by rCST is corroborated by the enhancement of small-world properties in the micro-level neural network (Table 3).

**Table 3.**
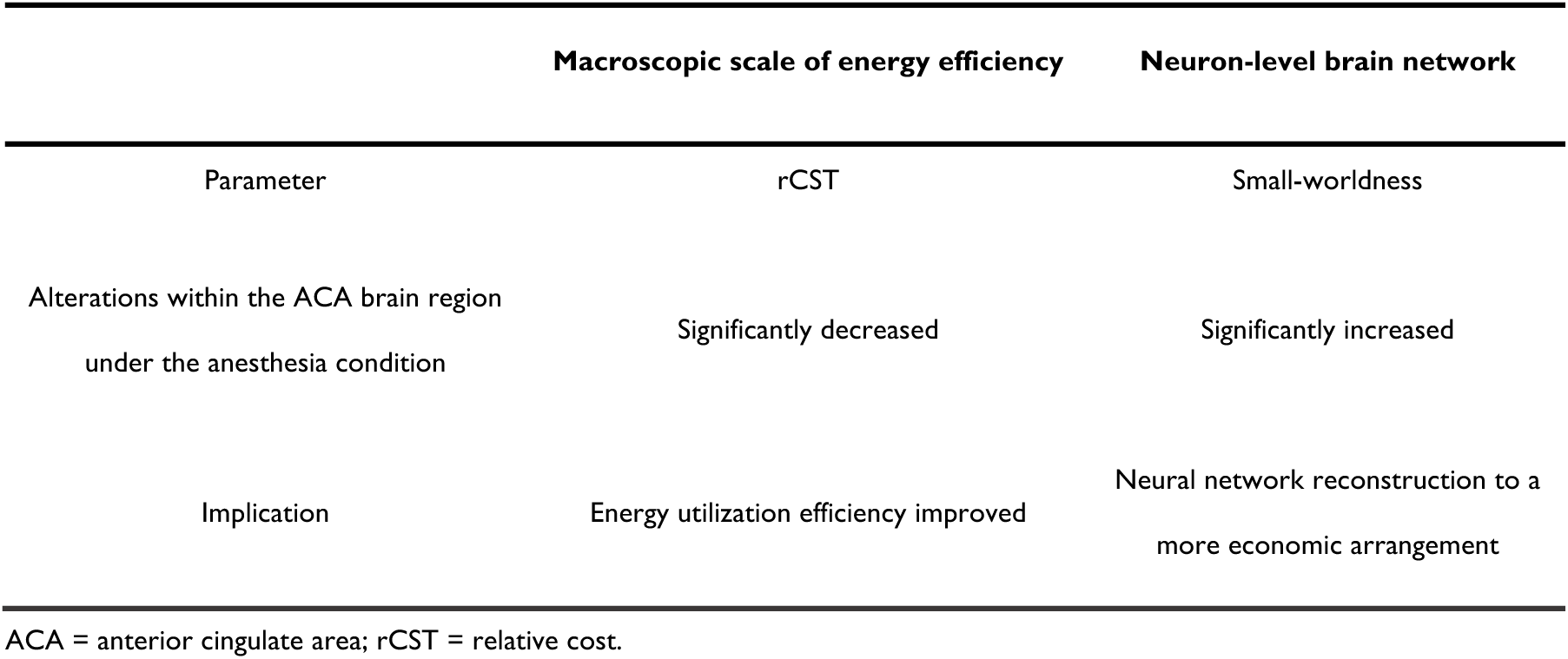
Macroscopic scale of energy efficiency correspondence to neuron-level brain network reconstruction.

## Discussion

Our research has revealed an intriguing similarity in the distribution of rPWR and rCST between humans and mice. Furthermore, compared with human studies, our mouse model has the ability to explore the rebalancing of energy utilization and the neuronal networks that underly this rebalancing. Using this, we have found a relationship between greater efficiency of energy utilization and the reconstruction of neuronal networks into a more economical form. We have also demonstrated that the brain’s economy reorganizes at the macroscopic and microscopic levels in response to anesthesia. As the macroscopic level can be imaged non-invasively in human patients through rPWR and rCST, this suggests that application of such medical imaging methods can locate the small, hard-to-detect changes which precede larger but irreversible changes in neurological disease.

Combining measurements of brain function and metabolism is very important to the study of neurodegeneration in disease. For example, metabolic and activity abnormalities in the ACA are linked to diseases like schizophrenia, where glucose metabolism decreases more than in other regions^62^, and neuropathic headaches^63^. Calibrated fMRI, a precise method for assessing brain function via CMR_O2_ quantification, traditionally utilizes gas challenges (administration of exogenous CO_2_ or O_2_) to elicit hypercapnia and modulate CMR_O2_ ^64, 65^. Herein, we employed a novel, gas-challenge-free, relaxometry-based calibrated fMRI method which has previously been used to track metabolic CMR_O2_ changes under anesthesia^35^ and to contrast oxygen versus glucose metabolism in the temporal lobes of healthy individuals and epilepsy patients^34^. This approach, while effective, is constrained by lengthy scanning sessions and stringent field homogeneity requirements. However, for the purposes of calculating rPWR and rCST, it allows measurement within a single imaging session on an MRI-only scanner. Concurrently, research by Castrillon et al. has explored the energetic implications of brain function by correlating glucose metabolism with connectivity, highlighting increased energy usage in fronto-parietal networks^66^. Unraveling the relationship between metabolic energy and neuronal activity is pivotal for brain function research.

Our results suggest that the rPWR and rCST method is highly sensitive and can be a powerful tool for potential research on brain diseases related to energy metabolism. Alzheimer’s Disease, characterized by glucose metabolism and mitochondrial dysfunction, is increasingly recognized as a metabolic disorder^67^. Studies have consistently reported reduced brain glucose metabolism in Alzheimer’s Disease, particularly in the anterior and posterior default mode network (DMN)^68^. Resting-state fMRI has been used to observe connectivity decline in the DMN and sensorimotor networks^69^. Additionally, research has indicated decreased DMN activity in mild cognitive impairment^70^, implying there may be reduced rPWR. The diminished small-worldness in the medial temporal lobe of AD patients^71^ suggests a potential increase in rCST. Given that Alzheimer’s Disease exhibits changes in both regional brain metabolism and activity^72, 73^, rPWR and rCST hold significant potential as biomarkers for diagnosis and measuring the progression of Alzheimer’s Disease. In the context of schizophrenia, impaired frontal cortex metabolism is a key pathogenic feature. Studies in mouse models have shown a reduction in glucose metabolism alongside heightened frontal cortex activation. Graph theoretical analyses point to an inefficient prefrontal network in schizophrenia, characterized by a lack of evenly distributed and highly clustered nodes^74^, which is an intriguing target to investigate as it may reflect an elevated rCST. This highlights the potential of rPWR and rCST as a metric for understanding the disorder’s neural underpinnings.

Our study demonstrates that rPWR and rCST are sensitive biomarkers of physiological state changes, exhibiting regionally heterogeneous alterations that reflect underlying neural network reconfigurations. Our findings also significantly contribute to the scientific and clinical understanding of brain function and metabolism, presenting novel avenues for research in these domains.

## Data Availability

Source data are provided with this paper. All data are available upon reasonable request.

## Acknowledgments

The authors would like to thank the 9.4 T Small Animal MRI Platform of Shanghai University of Science and Technology. We thank Xueying Fang and Songhai Xiong for their help with the 2PCI data analysis.

## Funding

ShanghaiTech University and the Shanghai Municipal Government (NZ and GJT)

National Natural Science Foundation of China Grant 81950410637 (GJT), National Natural Science Foundation of China Grant 3210055 (HL)

National Natural Science Foundation of China Grant 81771821 (ZL)

National Natural Science Foundation of China Grant 32170959 (NZ)

Strategic Priority Research Program of the Chinese Academy of Sciences Grant XDB32030100 (ZL)

Shanghai Municipal Science and Technology Major Project Grant 2018SHZDZX05 (ZL)

CAS Pioneer Hundred Talents Program (ZL).

## Author contributions

G.J.T. initiated, managed and supervised the project. G.J.T., and H.L. conceived and designed the experiments. D.W., H.L., Y.Z., J.G., M.X., B.B., M.P., performed the majority of experiments and data analysis. D.W., H.L., Y.Z., J.G., M.X., and N.Z. contributed to data analysis and interpretation. Z.L., N.Z., and G.J.T. supervised work. D.W., H.L., and G.J.T. created final figures and wrote the manuscript with contributions from all of the authors.

## Supplemental information

Figures S1-Figures S6

Table S1-S4

