## Supplementary figures and images for "Macroscopic cerebral energy efficiency corresponds to neuron reorganization in awake and anesthetized mice"

### Supplemental Fig. 1

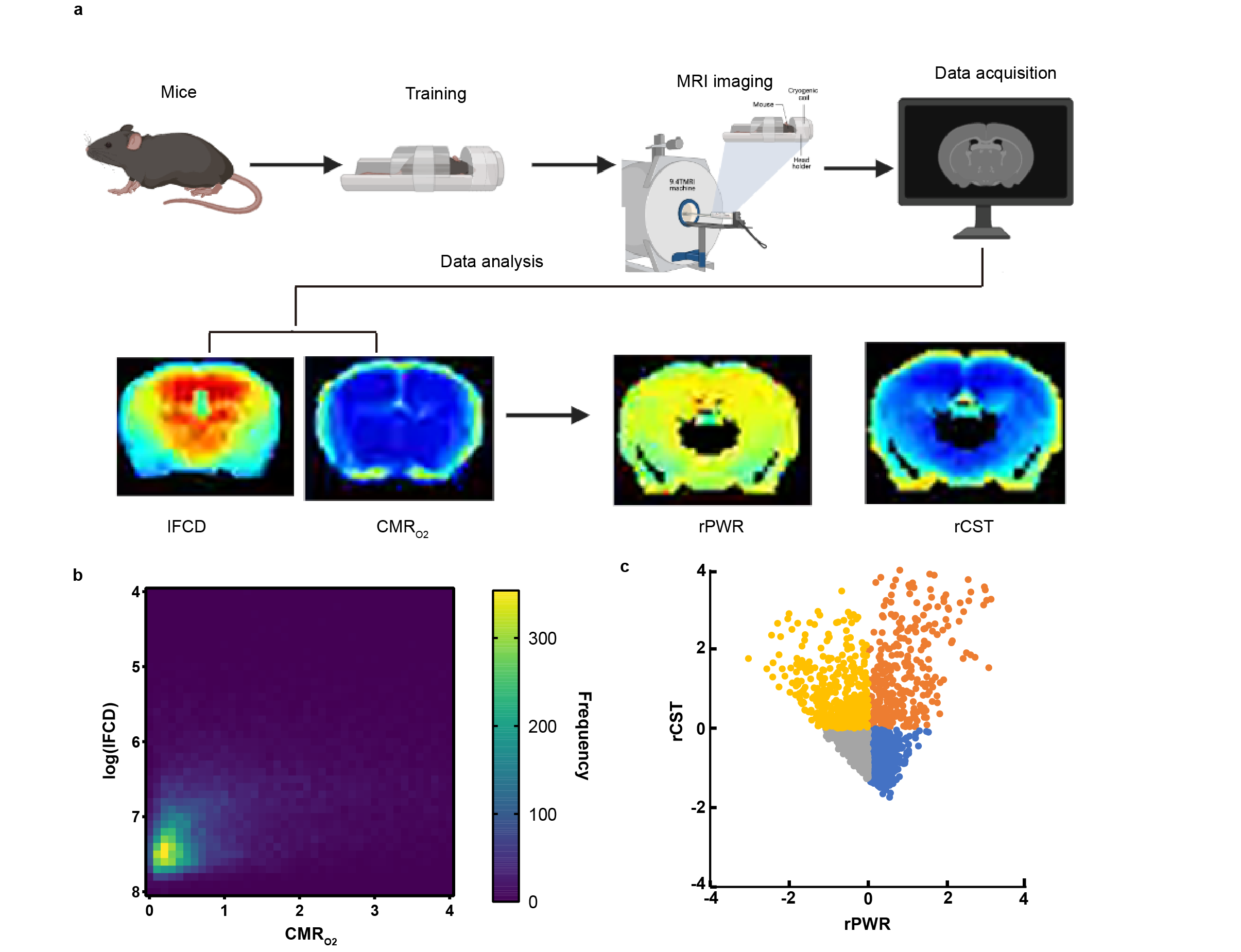

### Supplemental Fig. 2

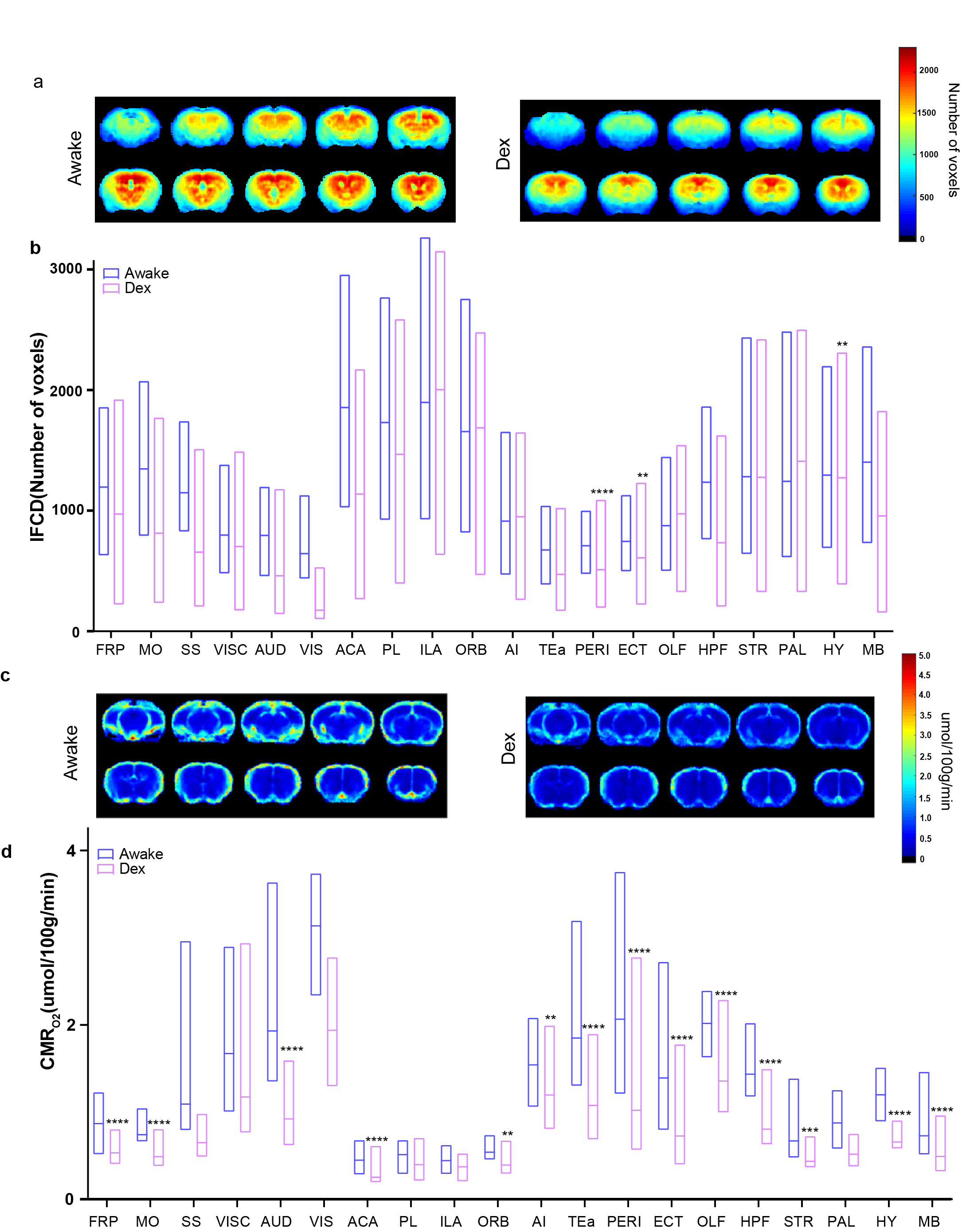

### Supplemental Fig. 3

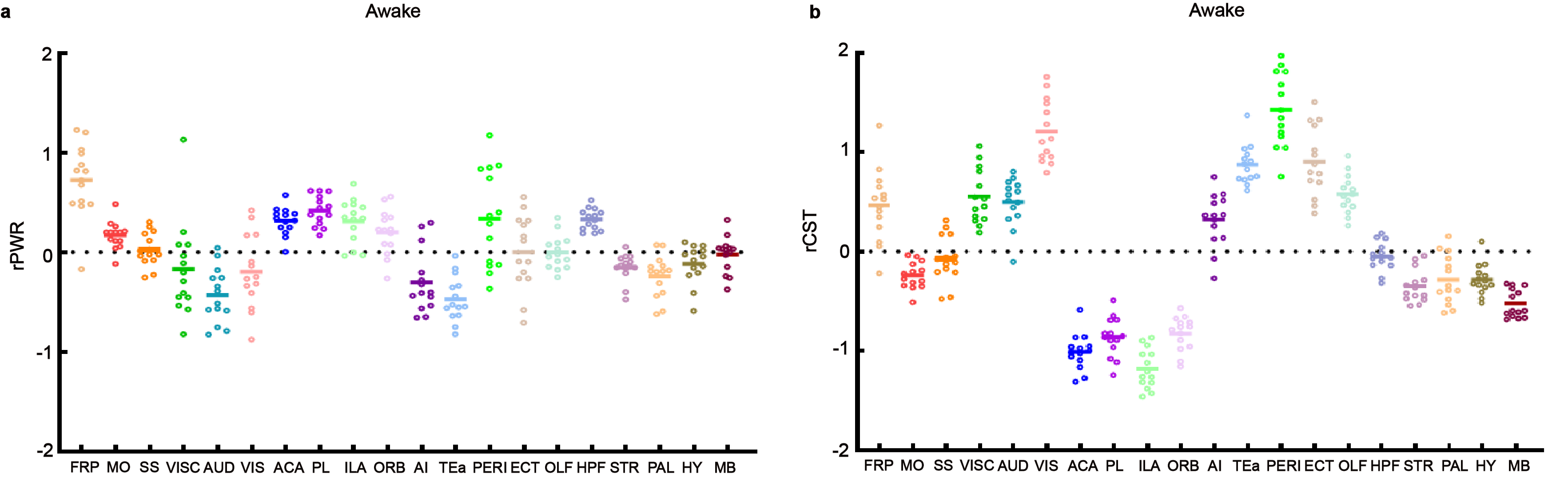

### Supplemental Fig. 4

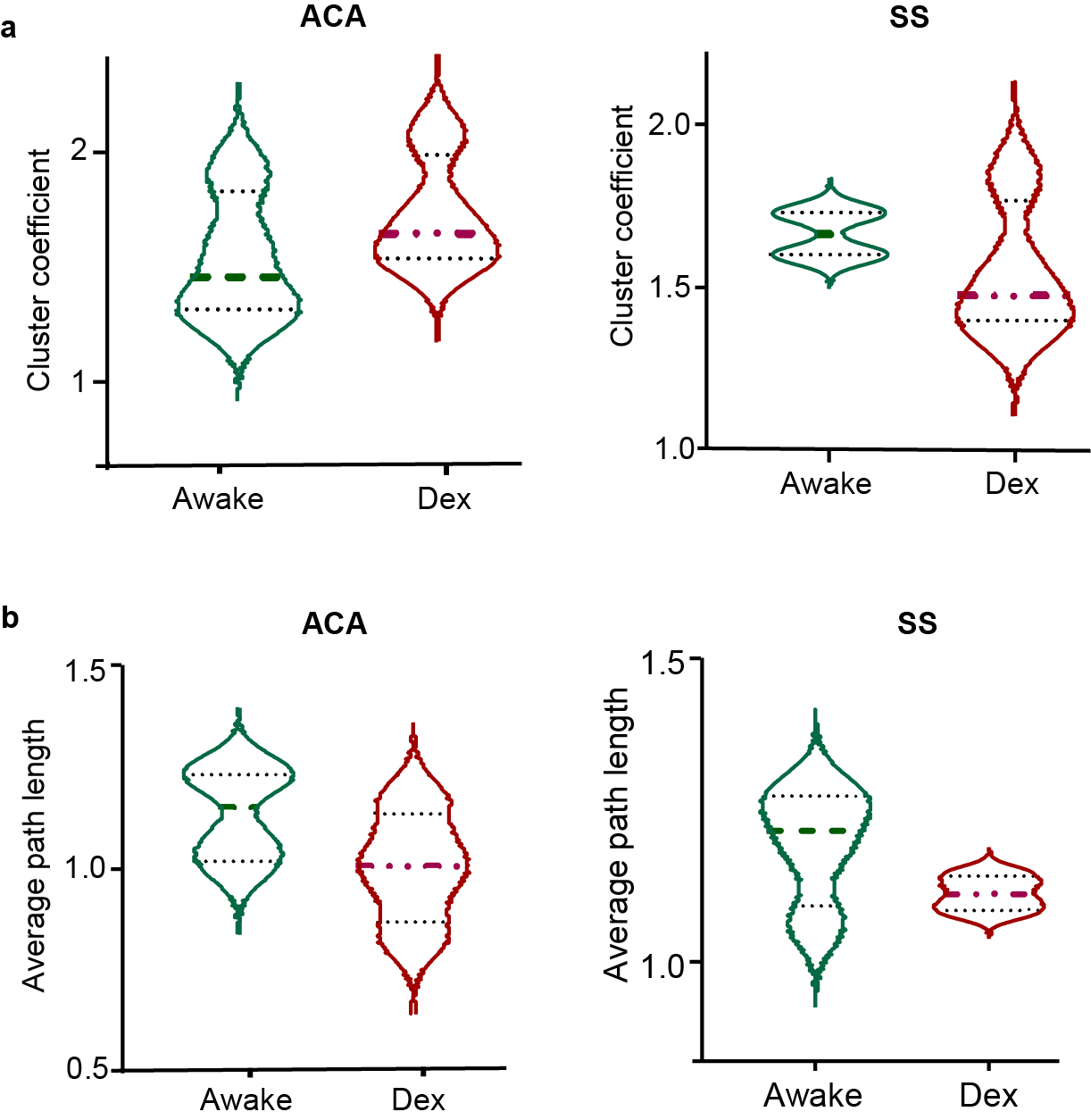

### Supplemental Fig. 5a

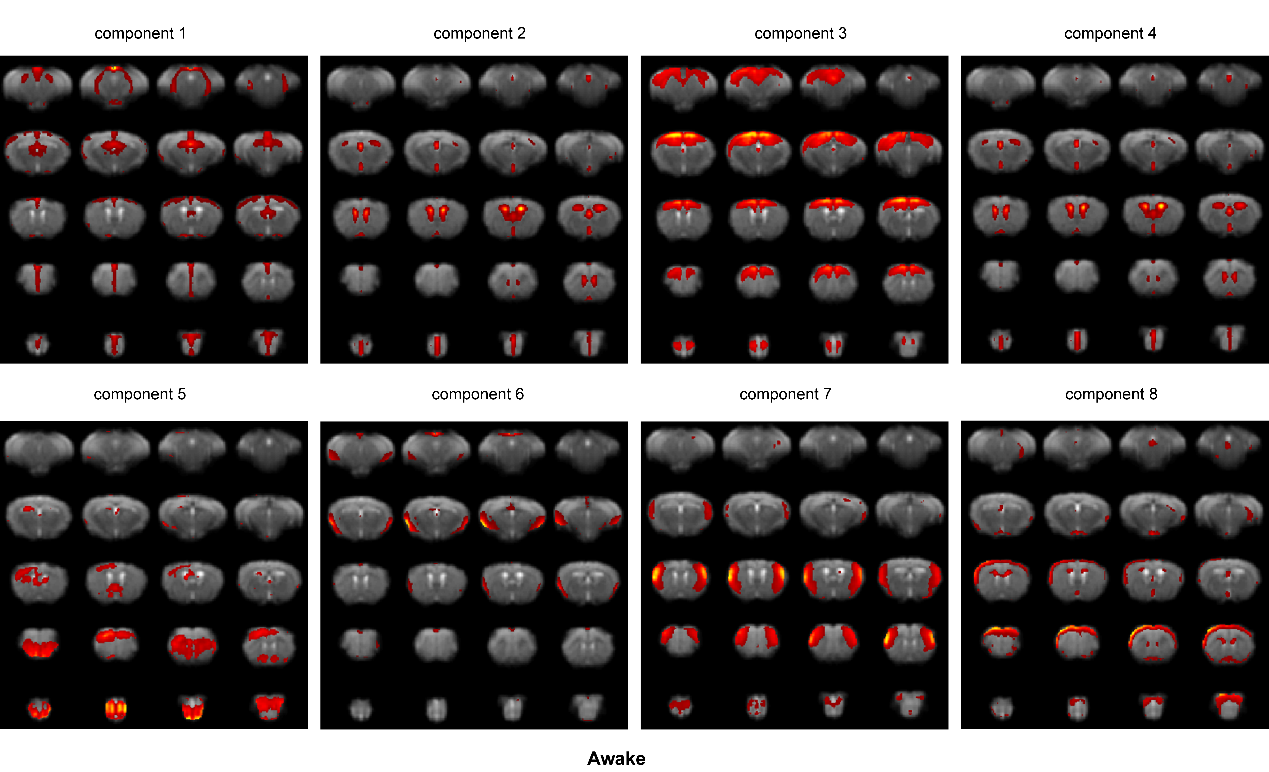

### Supplemental Fig. 5b

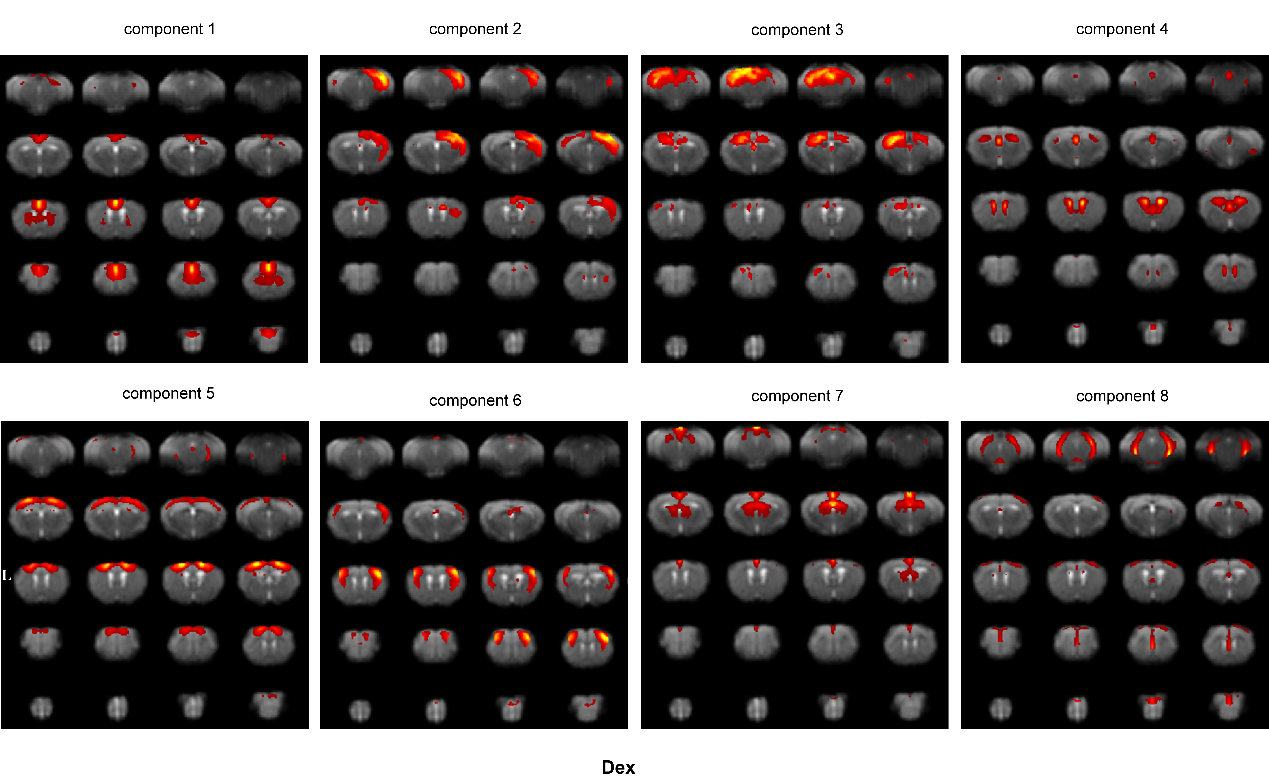

### Supplemental Fig. 6

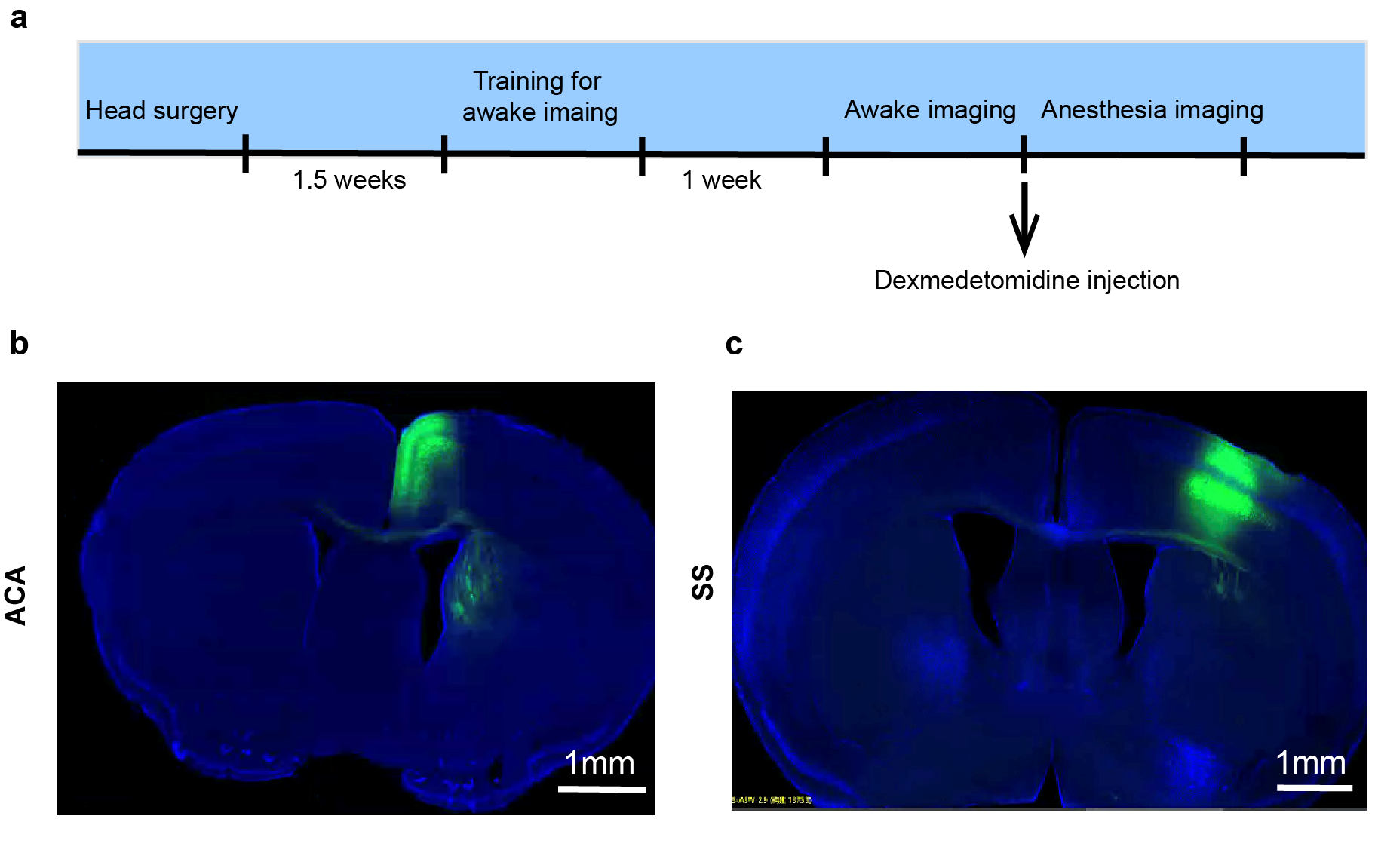
