## Supplemental Table. 1 for "Macroscopic cerebral energy efficiency corresponds to neuron reorganization in awake and anesthetized mice"

**Supplemental Table 1: List of brain region.**

| Abbreviation | Full name |
| --- | --- |
| ACA | Anterior cingulate area |
| AI | Agranular insular area |
| AUD | Auditory areas |
| CTXsp | Cortical subplate |
| ECT | Ectorhinal area |
| FRP | Frontal pole, cerebral cortex |
| GU | Gustatory areas |
| HPF | hippocampus formation |
| HY | hypothalamus |
| ILA | Infralimbic area |
| MB | midbrain |
| MO | Somatomotor areas |
| OLF | olfactory area |
| ORB | Orbital area |
| PAL | Pallidum |
| PERI | Perirhinal area |
| PL | Prelimbic area |
| PTLp | Posterior parietal association areas |
| RSP | Retrosplenial area |
| SS | Somatosensory areas |
| STR | Striatum |
| TE | Echo time |
| TEa | Temporal association areas |
| TH | thalamus |
| VIS | Visual areas |
| VISC | Visceral area |
