## Supplemental Table. 2 for "Macroscopic cerebral energy efficiency corresponds to neuron reorganization in awake and anesthetized mice"

**Supplemental Table 2: Results of statistically significant paired-*t* tests of rPWR, rCST, lFCD, and CMR_O2_.**

| Brain region | rPWR | | rCST | | lFCD | | CMR_O2_ | |
| --- | --- | --- | --- | --- | --- | --- | --- | --- |
|  | *p value* | *t* vlaue | *p value* | *t* vlaue | *p value* | *t* vlaue | *p value* | *t* vlaue |
| FRP | 1.10x10^-5^ | -5.42 | 2.38x10^-7^ | -6.91 |  |  | 1.72x10^-5^ | -6.60 |
| MO | 3.25x10^-2^ | -2.27 |  |  |  |  | 7.15x10^-7^ | -8.86 |
| VISC |  |  | 1.67x10^-6^ | 6.15 |  |  |  |  |
| AUD |  |  |  |  |  |  | 2.98x10^-6^ | -7.80 |
| VIS |  |  |  |  |  |  | 5.96x10^-7^ | -9.04 |
| ACA | 1.19x10^-7^ | 7.22 | 8.29x10^-4^ | 3.78 |  |  | 2.22x10^-4^ | -5.05 |
| PL |  |  | 4.11x10^-4^ | -4.05 |  |  | 1.29x10^-3^ | -4.08 |
| ORB | 3.15x10^-2^ | -2.27 | 1.79x10^-6^ | -6.12 |  |  | 1.57x10^-3^ | -3.98 |
| AI |  |  |  |  |  |  | 1.72x10^-3^ | -3.93 |
| TEa |  |  | 1.29x10^-4^ | 4.49 |  |  | 2.19x10^-5^ | -6.44 |
| PERI | 1.24 x10^-3^ | -3.62 | 1.66x10^-2^ | 2.56 | 9.87E-05 | 5.52 | 5.19x10^-6^ | -7.40 |
| ECT |  |  | 1.02x10^-4^ | 4.58 | 8.59E-03 | 4.3 | 2.22x10^-5^ | -6.43 |
| OLF | 1.45 x10^-2^ | 2.62 |  |  |  |  | 5.96x10^-8^ | -10.66 |
| HPF |  |  |  |  |  |  | 1.79x10^-7^ | -9.97 |
| CTXsp |  |  |  |  | 1.94x10^-3^ | 5.13 |  |  |
| STR | 1.22 x10^-2^ | -2.69 |  |  |  |  | 7.24x10^-4^ | -4.40 |
| PAL | 3.26 x10^-3^ | -3.24 |  |  |  |  | 2.32x10^-6^ | -7.98 |
| HY | 4.52 x10^-3^ | -3.11 |  |  | 0.01 | 4.14 | 5.96 x10^-8^ | -11.65 |
| MB |  |  |  |  |  |  | 1.35x10^-4^ | -5.34 |
