## Supplemental Table. 3 for "Macroscopic cerebral energy efficiency corresponds to neuron reorganization in awake and anesthetized mice"

**Supplemental Table 3:** **Result of the correlation between tSNR and lFCD, CMR_O2_, rPWR, and rCST from mice in Awake state (n=14), paired *t*-tests.** Values highlighted in yellow indicate brain regions with statistically significant correlation after multiple comparison correction (SGoF).

| Brain region | lFCD | CMR_O2_ | rPWR | rCST | Mean fMRI tSNR |
| --- | --- | --- | --- | --- | --- |
| FRP | 0.16 | -0.31 | -0.02 | -0.21 | 4.05 |
| MO | 0.14 | 0.06 | -0.08 | -0.25 | 3.32 |
| SS | 0.14 | 0.20 | 0.40 | -0.18 | 3.06 |
| GU | -0.52 | -0.80 | -0.72 | -0.61 | 2.88 |
| VISC | -0.20 | -0.52 | -0.54 | -0.30 | 2.50 |
| AUD | 0.07 | -0.32 | -0.13 | -0.26 | 3.14 |
| VIS | 0.16 | -0.33 | -0.42 | -0.11 | 1.99 |
| ACA | 0.50 | 0.12 | 0.13 | -0.02 | 7.46 |
| PL | 0.39 | -0.49 | -0.19 | -0.30 | 8.10 |
| ILA | 0.06 | -0.06 | 0.10 | 0.03 | 13.14 |
| ORB | -0.16 | -0.33 | 0.09 | -0.01 | 7.42 |
| AI | -0.30 | -0.43 | 0.04 | -0.02 | 2.69 |
| RSP | 0.58 | -0.37 | -0.25 | -0.50 | 2.06 |
| PTLp | 0.50 | -0.80 | -0.61 | -0.77 | 2.03 |
| TEa | -0.09 | -0.41 | -0.24 | -0.43 | 2.14 |
| PERI | -0.29 | -0.41 | -0.17 | -0.40 | 2.05 |
| ECT | -0.28 | -0.41 | -0.32 | -0.27 | 2.44 |
| OLF | -0.18 | -0.30 | 0.13 | -0.10 | 1.60 |
| HPF | -0.43 | -0.38 | 0.04 | 0.18 | 1.84 |
| CTXsp | -0.45 | -0.72 | -0.47 | -0.08 | 2.02 |
| STR | 0.09 | -0.56 | -0.04 | -0.48 | 4.46 |
| PAL | 0.22 | -0.42 | 0.30 | -0.59 | 6.54 |
| TH | -0.17 | -0.72 | 0.22 | -0.22 | 8.68 |
| HY | 0.24 | -0.70 | -0.64 | -0.48 | 6.18 |
| MB | -0.13 | -0.37 | -0.09 | 0.10 | 5.08 |
| mean | 0.26 | 0.18 | 0.24 | 0.19 | 2.29 |
| std | 0.31 | 0.28 | 0.31 | 0.25 | 2.88 |
| p_value | 0.00 | 0.00 | 0.07 | 0.00 |  |
