## Supplemental Table. 4 for "Macroscopic cerebral energy efficiency corresponds to neuron reorganization in awake and anesthetized mice"

**Supplemental Table 4: Result of the correlation between tSNR and lFCD, CMR_O2_,rPWR and rCST from mice in Dex state ( n=14 ), paired *t*-test.** Values highlighted in yellow indicate brain regions with statistically significant correlation after multiple comparison ( SGoF ).

| Brain region | lFCD | CMR_O2_ | rPWR | rCST | Mean fMRI tSNR |
| --- | --- | --- | --- | --- | --- |
| FRP | 0.15 | 0.53 | 0.32 | 0.05 | 4.09 |
| MO | -0.20 | -0.43 | 0.41 | -0.63 | 3.07 |
| SS | -0.14 | -0.24 | -0.02 | 0.14 | 2.83 |
| GU | 0.13 | -0.10 | 0.07 | -0.03 | 2.90 |
| VISC | 0.24 | -0.09 | -0.16 | 0.01 | 2.64 |
| AUD | -0.17 | -0.12 | -0.16 | -0.24 | 3.27 |
| VIS | -0.17 | -0.08 | -0.09 | -0.28 | 2.09 |
| ACA | -0.28 | -0.08 | 0.06 | 0.18 | 7.36 |
| PL | -0.49 | 0.05 | 0.59 | -0.34 | 9.47 |
| ILA | -0.20 | 0.09 | -0.01 | 0.14 | 12.65 |
| ORB | -0.24 | -0.08 | 0.38 | -0.14 | 6.93 |
| AI | 0.15 | 0.14 | 0.12 | 0.22 | 3.02 |
| RSP | -0.04 | 0.70 | 0.40 | 0.56 | 2.08 |
| PTLp | 0.08 | 0.49 | 0.33 | 0.56 | 2.15 |
| TEa | -0.47 | -0.24 | 0.03 | -0.45 | 2.24 |
| PERI | -0.12 | -0.31 | 0.13 | -0.44 | 2.22 |
| ECT | -0.30 | -0.41 | 0.24 | -0.55 | 2.54 |
| OLF | -0.26 | 0.56 | 0.21 | 0.49 | 1.61 |
| HPF | -0.57 | -0.19 | -0.39 | -0.23 | 1.91 |
| CTXsp | -0.42 | 0.36 | 0.63 | -0.26 | 2.25 |
| STR | -0.07 | 0.04 | -0.22 | 0.16 | 4.28 |
| PAL | -0.06 | -0.14 | -0.12 | 0.05 | 6.54 |
| TH | -0.21 | -0.46 | -0.12 | 0.13 | 8.20 |
| HY | 0.58 | 0.19 | -0.17 | -0.11 | 4.36 |
| MB | 0.02 | 0.18 | -0.24 | 0.24 | 4.77 |
| mean | 0.19 | 0.25 | 0.22 | 0.27 | 2.13 |
| std | 0.26 | 0.31 | 0.26 | 0.33 | 2.79 |
| p_value | 0.00 | 0.00 | 0.03 | 0.00 |  |
